# Forskolin as a multi-target modulator of the oncogenic PI3K/Akt signaling pathway: An *in silico* integrative GO/KEGG enrichment analysis and network pharmacology

**DOI:** 10.1101/2025.05.11.653344

**Authors:** Eman M. Sarhan

## Abstract

Forskolin (FSK), a natural diterpene plant metabolite, exhibits promising anticancer properties, yet its molecular mechanisms remain to be fully elucidated. The current study employs an *in silico* integrative framework to systematically evaluate FSK anticancer mechanistic potential, paving its repurposing from a traditional natural remedy to a validated, multitargeted oncological therapy. Database mining was used to predict cancer and FSK-related protein targets. Then, the intersect FSK/Cancer common core targets [n=89] were identified. A protein-protein interaction (PPI) network was constructed to identify the gene hubs of core targets. Functional enriched GO/KEGG annotation and cluster analysis were performed to predict molecular pathways and metabolic modules involved in FSK anti-cancer effect. Our results reveal that FSK predominantly targets the PI3K/Akt signaling pathway and its associated oncogenic network. Significant enrichment in PI3K/Akt, EGFR, and Ras pathways, alongside key metabolic modules, underscores FSK potential to inhibit proliferation, induce apoptosis, and alter cancer metabolism. Among the 14 hub genes identified, AKT1, EGFR, PIK3CA, and TP53 further highlight FSK multi-targeted mechanism. These findings provide a molecular framework for the continued development of FSK as a targeted anticancer therapy.

## INTRODUCTION

Forskolin (C_22_H_34_O_7_; Mwt. 410.5 g/mol), also commonly known as colforsin, coleonol, and colforsina, is a natural labdane diterpene plant metabolite derived from the roots (Bhat et al., 1977; Miastkowska et al., 2017; Salehi et al., 2019; Singh and Tandon, 1982; Tripathi et al., 1995) and stems (T Pullaiah, 2022; Roshni and Rekha, 2024; Saleem et al., 2006) of *Coleus forskohlii* (Willd.) Briq. (Lamiaceae). *C. forskohlii*, which has been commonly used in traditional Indian Ayurvedic medicine and South Asian subtropical regions (T. Pullaiah, 2022; Salehi et al., 2019) is considered to be the most common, if not the only known natural source of forskolin (Roshni and Rekha, 2024). Forskolin has been used for centuries in traditional medicine and is well-documented for its safety and efficacy in modern medical studies (Genovese et al., 2018; Godard et al., 2005; Henderson et al., 2005; Illiano et al., 2018; Loftus et al., 2015; Pateraki et al., 2017). It has been widely recognized for its diverse pharmacological properties, including its therapeutic potential for colic, painful urination, eczema, convulsions, insomnia, hypertension, congestive heart failure, and respiratory disorders (Lokesh et al., 2018; T. Pullaiah, 2022). It is the only plant-derived compound known to be a rapid and reversible activator of the adenylate cyclase enzyme, subsequently elevating the intracellular cyclic adenosine monophosphate (cAMP) levels—a secondary messenger integral to modulating diverse cellular processes and disease pathogenesis, as well as impacting many drugs’ mechanisms of action (T. Pullaiah, 2022).

Beyond its historical applications, modern research has unveiled forskolin’s multifaceted therapeutic repurposing, particularly in oncology, where its ability to disrupt oncogenic signaling pathways and enhance chemosensitivity of conventional drugs has garnered significant attention (Illiano et al., 2018; Wang et al., 2019). Many preclinical investigations have revealed forskolin’s anticancer efficacy across diverse malignancies including glioblastoma (GBM) (He et al., 2025), multiple myeloma (MM) (Follin-Arbelet et al., 2015), non-Hodgkin’s lymphomas (NHLs) (Wang et al., 2019), acute myeloid leukemia (AML) (Illiano et al., 2018), breast (Hayashi et al., 2018), bancreatic (Quinn et al., 2017), gastric (Li and Wang, 2006) cancer types. However, the exact molecular mechanisms underlying forskolin’s anticancer activity remain incompletely defined. Traditional *in vitro* and *in vivo* investigations, while invaluable, often lack the precision necessary to map dynamic protein-ligand interactions or predict off-target consequences comprehensively. This gap emphasizes the critical role of *in silico* multi-omics integrative analysis to identify therapeutic targets, validate binding affinities, and unravel molecular-level mechanisms.

The current study employs an *in silico* integrative framework to systematically evaluate forskolin’s anticancer mechanistic potential, paving its repurposing from a traditional natural remedy to a validated, multitargeted oncological therapy.

## MATERIALS AND METHODS

### FSK-target prediction

FSK-related protein targets have been retrieved using the STITCH database (http://stitch.embl.de/, accessed on 3 May 2025), Super-PRED (https://prediction.charite.de/index.php, accessed on 3 May 2025), and PharmMapper (http://www.lilab-ecust.cn/pharmmapper/, accessed on 7 May 2025) (Liu et al., 2010; Wang et al., 2017, 2016) servers. AnnotationDbi v1.66.0 with the human annotation database org.Hs.eg.db v3.19.1 R packages were used for mapping the different gene identifiers retrieved from Super-PRED and PharmMapper. Then, all deduplicated FSK-related targets were gathered to construct a single FSK dataset.

### Cancer-target prediction

Cancer-related targets were retrieved using text mining of the searched keywords ‘Cancer’, ‘Carcinoma’, and ‘Carcinogenesis’ from the DisGeNET (https://disgenet.com/, accessed on 3 May 2025), Disease (https://diseases.jensenlab.org/, accessed on 3 May 2025), and GeneCards (https://www.genecards.org/, accessed on 3 May 2025) databases. The top 250 highly scored targets of each keyword dataset were filtered individually, and then the deduplicated records were combined to construct cancer datasets.

### Protein-protein interaction (PPI) network construction and analysis

The common targets’ intersection between FSK and cancer datasets has been visualized using the VennDiagram v1.7.3 R package. The PPI network of FSK/Cancer core targets was constructed using the STRING v12.0 (https://string-db.org/, accessed on 8 May 2025) database. A list of 89 input SYMBOL identifiers was analyzed at a high confidence score > 0.7 using *Homo sapiens* annotation species. The network constructed was then analyzed and visualized using Cytoscape v3.10.3 software to identify the hub genes.

### Gene ontology (GO) and Kyoto Encyclopedia of Genes and Genomes (KEGG) enrichment analysis

Prior to the functional enrichment analysis, the AnnotationDbi v1.66.0 R package was used to map the core targets’ SYMBOL gene identifiers into ENTREZID. Gene ontology (GO) as *well as* Kyoto Encyclopedia of Genes and Genomes (KEGG) pathways and metabolic modules were then analyzed using the *clusterProfiler* v4.12.6 R package (https://bioconductor.org/packages/release/bioc/html/clusterProfiler.html). GO terms of biological processes (BP), cellular components (CC), and molecular functions (MF) were investigated using *enrichGO* function with the method of Benjamini-Hochberg correction (BH) at pvalue < 0.05, p.adjust < 0.01, and qvalue < 0.05. KEGG pathways were identified using *enrichKEGG* function at fold enrichment (FDR) pvalue < 0.05, p.adjust < 0.05, and qvalue < 0.05, while those metabolic modules were identified using *enrichMKEGG* function. The generated data were ascendingly sorted by p.adjust and then descendingly by FoldEnrichment to identify the most significant terms, pathways, and modules. Top 5 enriched results were visualized by enrichment plots and gene-concept network graphs using ggplot2 v3.5.1 and enrichplot v1.24.4 R packages, respectively. The pathway gene expression map was visualized using the browseKEGG function.

## RESULTS AND DISCUSSION

### FSK/Cancer-related target prediction

Cancer datasets retrieved from *DisGeNET*, Disease, and GeneCards databases were sorted by scoreGDA, disease score, and relevance score, respectively. The top 250 highly scored records were filtered from each dataset individually and then merged into a combined cancer dataset with a total of 716 deduplicated target records (Figure 1A). On the other hand, all FSK-related targets gathered from three different databases (Super-PRED, PharmMapper, and STITCH) were combined after removing irrelevant, empty, and duplicated records to form a single FSK dataset with a total of 506 target records (Figure 1B). FSK/Cancer intersect [n = 89] represents the core common targets with a probable FSK anticancer potential (Figure 1C).

**Fig. 1:**
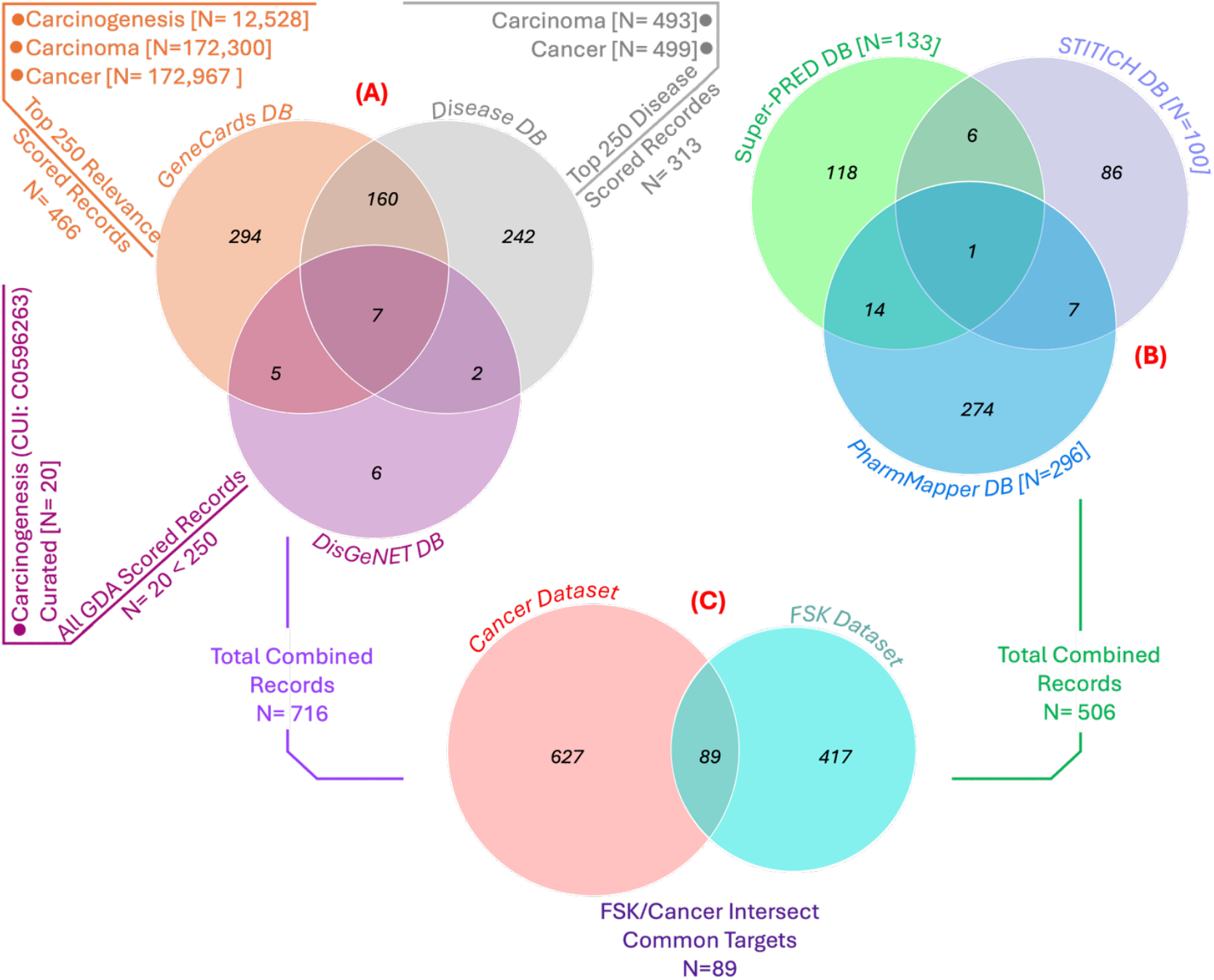
FSK/Cancer Core targets prediction workflow. *Venn diagrams represent (A) Cancer datasets retrieved from DisGeNET (Magenta), GeneCards (Orange), and Disease (Grey) databases; (B) FSK datasets retrieved from Super-PRED (Green), STITCH (Violet), and PharmMapper (Deepskyblue) databases; (C)* FSK/Cancer *core common targets intersect [n=89].*

### Protein-protein interaction (PPI) network construction and analysis

The PPI network of FSK/Cancer core targets constructed using the STRING database at a high confidence score > 0.7 and average node degree > 17.2 generates 88 nodes connected with 765 edges (Figure 2A). Fourteen hub genes (Figure 2B) of the highly connected nodes at a degree centrality >30 are supposed to represent the potential FSK therapeutic targets in cancer incidence and progression.

**Fig. 2:**
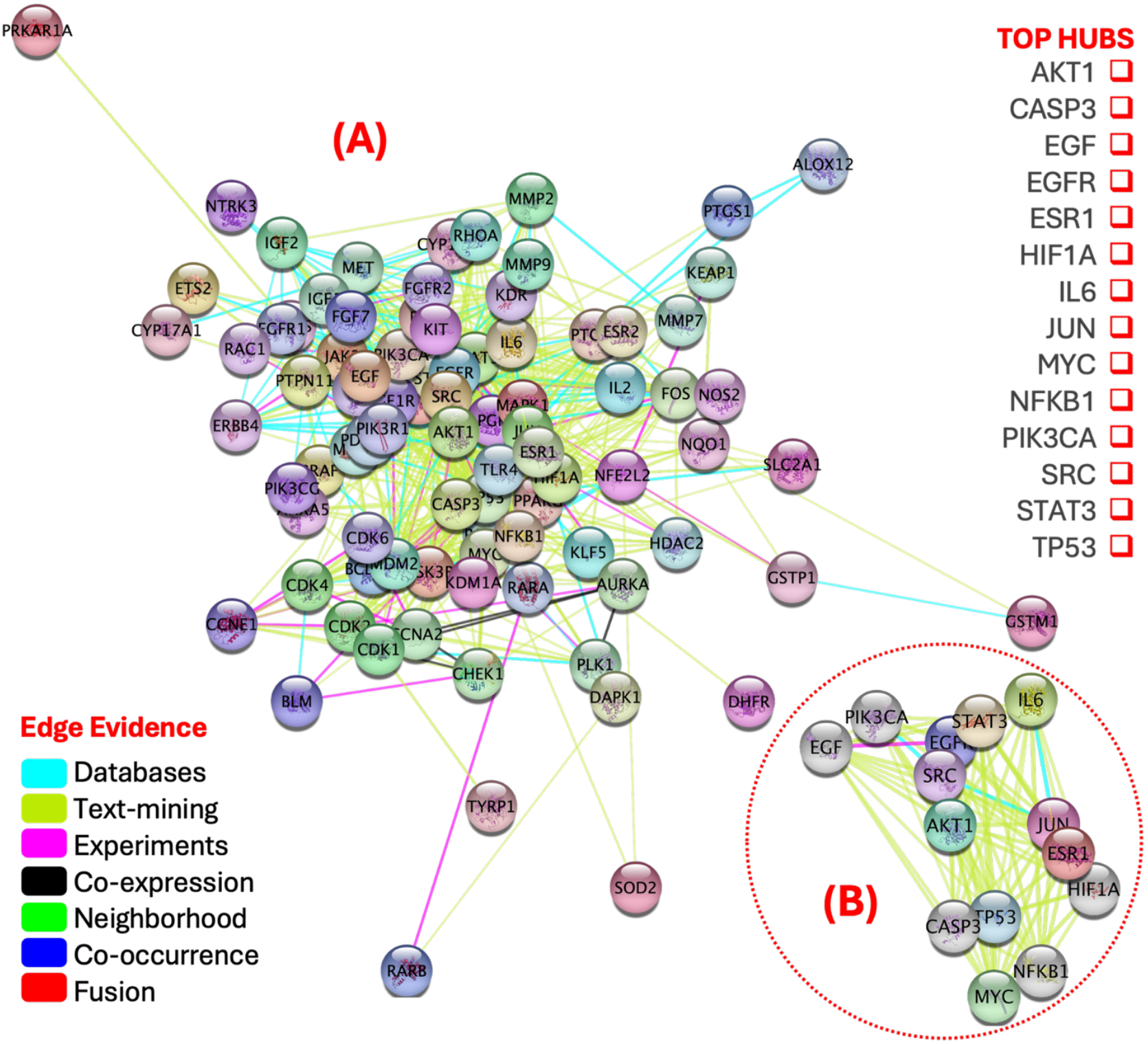
Protein-protein interaction (PPI) network of (A) FSK/Cancer core target genes (degree> 17.2) and (B) hub genes (n = 14; degree>30).

### GO/KEGG enrichment analysis

FSK demonstrates multifaceted anticancer modulation of oncogenic signaling pathways and molecular mechanisms as evidenced by its associated gene ontology (GO) terms and KEGG pathways and modules. GO and KEGG enriched analysis of FSK/Cancer core targets [N=89] was performed using the *clusterProfiler* R package and resulted in a total of 1646 GO terms [1526 BP, 31 CC, and 89 MF], 150 KEGG pathways, and 3 metabolic modules, of which the top significant terms, pathways, and modules are listed in Table 1-2 and Figure 3-4.

**Table 1:** Top 5 *BP, CC, and MF* gene ontology (GO) terms of FSK/Cancer common targets.

|  | ID | Description | FDR | pvalue | p.adjust | qvalue | N | Ont |
| --- | --- | --- | --- | --- | --- | --- | --- | --- |
| 1 | GO:0062197 | cellular response to chemical stress | 19.79 | 4.74E-30 | 1.73E-26 | 7.38E-27 | 29 | BP |
| 2 | GO:0006979 | response to oxidative stress | 16.00 | 2.50E-28 | 4.56E-25 | 1.94E-25 | 30 | BP |
| 3 | GO:0033002 | muscle cell proliferation | 21.98 | 4.31E-28 | 5.24E-25 | 2.23E-25 | 26 | BP |
| 4 | GO:0000302 | response to reactive oxygen species | 24.05 | 9.32E-26 | 8.51E-23 | 3.62E-23 | 23 | BP |
| 5 | GO:0034599 | cellular response to oxidative stress | 20.54 | 3.39E-25 | 2.48E-22 | 1.05E-22 | 24 | BP |
| 16 | GO:1902911 | protein kinase complex | 13.41 | 2.35E-08 | 2.81E-06 | 1.95E-06 | 9 | CC |
| 17 | GO:0061695 | transferase complex, transferring phosphorus-containing groups | 8.71 | 1.25E-08 | 2.81E-06 | 1.95E-06 | 12 | CC |
| 18 | GO:0000307 | cyclin-dependent protein kinase holoenzyme complex | 23.95 | 1.83E-07 | 1.10E-05 | 7.58E-06 | 6 | CC |
| 19 | GO:1902554 | serine/threonine protein kinase complex | 13.15 | 1.72E-07 | 1.10E-05 | 7.58E-06 | 8 | CC |
| 20 | GO:0044853 | plasma membrane raft | 13.85 | 7.42E-07 | 3.55E-05 | 2.45E-05 | 7 | CC |
| 31 | GO:0004714 | transmembrane receptor protein tyrosine kinase activity | 38.15 | 4.71E-15 | 9.04E-13 | 5.30E-13 | 11 | MF |
| 32 | GO:0004713 | protein tyrosine kinase activity | 21.11 | 3.93E-15 | 9.04E-13 | 5.30E-13 | 14 | MF |
| 33 | GO:0061629 | RNA polymerase II-specific DNA-binding transcription factor binding | 10.92 | 3.87E-14 | 4.95E-12 | 2.91E-12 | 18 | MF |
| 34 | GO:0140297 | DNA-binding transcription factor binding | 8.71 | 8.87E-14 | 8.51E-12 | 4.99E-12 | 20 | MF |
| 35 | GO:0019199 | transmembrane receptor protein kinase activity | 28.98 | 1.15E-13 | 8.86E-12 | 5.20E-12 | 11 | MF |

**Fig. 3:**
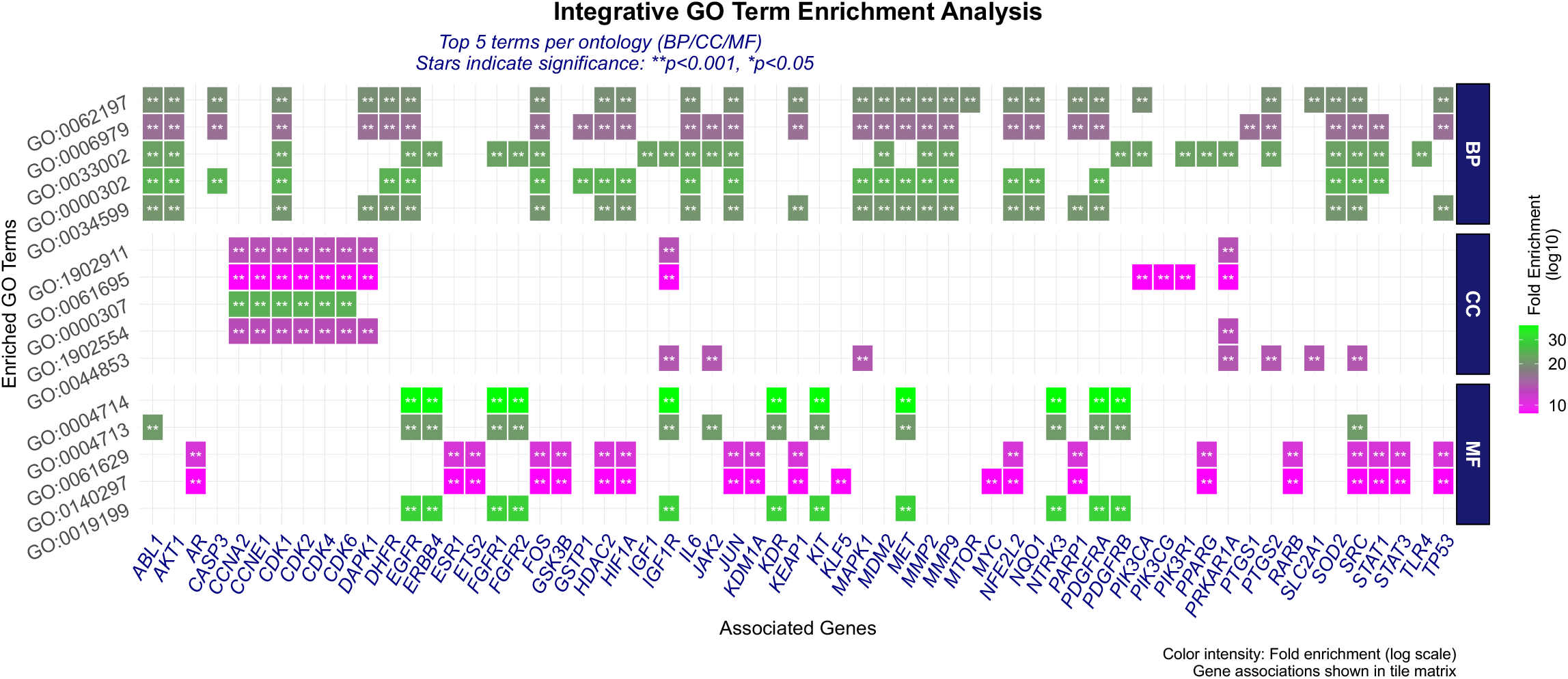
Heatmap of the top 5 *BP, CC, and MF* gene ontology (GO) terms of FSK/Cancer common targets. Associated protein targets *are expressed on the horizontal x-axis, while GO terms are expressed on the vertical y-axis according to p.adjust values. The color scale represents the fold enrichment*.

Our GO results provide valuable insights into the potential FSK stress-response mechanisms and targeted oncogenic pathways (Table 1; Figure 3). The biological processes (BP) associated with FSK anticancer activity highlight its potential in modulating cellular reactive oxygen species (ROS) (GO:0000302) and responding to chemical (GO:0062197; FDR: 19.79) and/or oxidative stress (GO:0006979; FDR: 16.00 and GO:0034599; FDR: 20.54), which are critical in cancer biology. FSK can modulate the oxidative stress (OS) and enhance the cellular antioxidant defense, reducing damage from reactive oxygen species (ROS). It can also modulate the expression of the forkhead transcription factor O1 (FOXO1) gene and its target gene catalase, essentially regulating metabolism and OS (Elwia et al., 2018). The term muscle cell proliferation (GO:0033002; FDR: 21.98) suggests that FSK may also influence muscle cell dynamics, potentially impacting tumor microenvironments where muscle tissue interacts with cancer cells.

The GO terms related to cellular components (CC) reveal that spatial regulation of kinase signaling and membrane domain organization are crucial in understanding FSK’s anticancer mechanisms. FSK targets kinase complexes (GO:1902911 with FDR of 13.41 and GO:1902554 with FDR of 13.15) critical for cell cycle regulation and cancer progression. It activates adenylate cyclase, increasing cAMP levels, which in turn activates protein kinase A (PKA) and affects downstream signaling pathways, including those involving protein kinase C (PKC) and other kinases (Illiano et al., 2018; Sapio et al., 2017). Studies demonstrate its inhibition of DEAD-box RNA helicase 3 (DDX3) and cyclooxygenase-2 (COX-2) expression in cervical cancer (Ravinder et al., 2022), disrupting RNA helicase-kinase interaction essential for tumor growth. FSK also modulates Axin/β-catenin signaling in NHLs by altering kinase-phosphatase equilibria, leading to β-catenin degradation and apoptosis (Wang et al., 2019). This aligns with findings that oncogenic kinases like PI3K/AKT and MAPK converge on β-catenin phosphorylation, causing its stabilization and nuclear translocation (Blume-Jensen and Hunter, 2001). FSK interferes with phosphate-transferase enzymes (GO:0061695; FDR: 8.71), including cyclin-dependent kinases (CDKs) and receptor tyrosine kinases (RTKs). Another significant term is the cyclin-dependent protein kinase holoenzyme complex (GO:0000307; FDR: 13.15) that reflects FSK’s ability to disrupt CDK-cyclin assemblies critical for cell cycle progression. It can also alter plasma membrane lipid raft (GO:0044853; FDR: 13.85) composition, influencing the localization and activity of kinases, which further impacts oncogenic cell signaling (Wu et al., 2003).

Moreover, the significant enrichment of FSK’s molecular function GO-terms, such as transmembrane receptor protein tyrosine kinase activity (GO:0004714; FDR: 38.15) and protein tyrosine kinase activity (GO:0004713; FDR: 21.11), suggests that FSK may modulate critical signaling cascades involved in cancer progression.

KEGG analysis of the FSK/Cancer common targets [N=89] revealed the probable FSK’s potential efficacy on multiple disease, signaling, and metabolic pathways (Table 2; Figure 4). The PI3K/Akt signaling pathway (hsa04151; FDR: 9.66; p-value: 6.55E-27; Count: 36) is the most significant enriched pathway, being a master regulator of cell growth, survival, metabolism, and apoptosis, and its dysregulation is a hallmark of many cancers (Manning and Toker, 2017; Saxton and Sabatini, 2017). Other top-enriched pathways, including EGFR tyrosine kinase inhibitor resistance (hsa01521; FDR: 25.49), the Ras signaling pathway (hsa04014; FDR: 9.79), and various cancer types (Figure 5), are tightly interconnected with PI3K/Akt, highlighting FSK’s potential to disrupt oncogenic signaling networks at multiple levels. Among the 36 genes engaging in the PI3K/Akt signaling (Figure 6), 8 are of the hub genes previously identified in Figure 2 with highly PPI-connected nodes (degree > 30), including AKT1, EGF, EGFR, IL6, MYC, NFKB1, PIK3CA, and TP53. Almost all of these hubs are engaged in the regulation of the 16 human cancer types illustrated in Figure 5.

**Table 2:** Top 15 KEGG pathways and metabolic modules of FSK/Cancer common targets.

| ID | Description | FDR | pvalue | p.adjust | qvalue | N | Cat |
| --- | --- | --- | --- | --- | --- | --- | --- |
| 1 hsa04151 | PI3K-Akt signaling pathway | 9.66 | 5.65E-27 | 1.30E-24 | 4.52E-25 | 36 | Pathway |
| 2 hsa05215 | Prostate cancer | 22.79 | 7.27E-26 | 5.60E-24 | 1.94E-24 | 23 | Pathway |
| 3 hsa05205 | Proteoglycans in cancer | 13.81 | 5.44E-26 | 5.60E-24 | 1.94E-24 | 29 | Pathway |
| 4 hsa01521 | EGFR tyrosine kinase inhibitor resistance | 25.49 | 9.14E-25 | 5.28E-23 | 1.83E-23 | 21 | Pathway |
| 5 hsa05224 | Breast cancer | 15.09 | 1.79E-21 | 8.29E-20 | 2.87E-20 | 23 | Pathway |
| 6 hsa01522 | Endocrine resistance | 19.62 | 4.15E-21 | 1.60E-19 | 5.53E-20 | 20 | Pathway |
| 7 hsa05218 | Melanoma | 23.95 | 9.53E-21 | 3.15E-19 | 1.09E-19 | 18 | Pathway |
| 8 hsa05230 | Central carbon metabolism in cancer | 23.25 | 2.16E-19 | 6.25E-18 | 2.16E-18 | 17 | Pathway |
| 9 hsa05167 | Kaposi sarcoma-associated herpesvirus infection | 11.40 | 1.27E-18 | 3.26E-17 | 1.13E-17 | 23 | Pathway |
| 10 hsa05161 | Hepatitis B | 12.51 | 7.01E-18 | 1.52E-16 | 5.28E-17 | 21 | Pathway |
| 11 hsa04014 | Ras signaling pathway | 9.79 | 7.26E-18 | 1.52E-16 | 5.28E-17 | 24 | Pathway |
| 12 hsa05225 | Hepatocellular carcinoma | 12.00 | 1.70E-17 | 3.28E-16 | 1.13E-16 | 21 | Pathway |
| 13 hsa05214 | Glioma | 20.44 | 2.54E-17 | 4.52E-16 | 1.57E-16 | 16 | Pathway |
| 14 hsa05212 | Pancreatic cancer | 20.18 | 3.19E-17 | 5.26E-16 | 1.82E-16 | 16 | Pathway |
| 15 hsa05162 | Measles | 13.27 | 1.02E-16 | 1.57E-15 | 5.44E-16 | 19 | Pathway |
| 16 M00977 | C19-Steroid hormone biosynthesis (androgen backdoor pathway),<br>pregnenolone => androsterone => dihydrotestosterone | 22.23 | 0.04 | 0.05 | 0.04 | 1 | Module |
| 17 M00034 | Methionine salvage pathway | 18.52 | 0.05 | 0.05 | 0.04 | 1 | Module |
| 18 M00976 | C19-Steroid hormone biosynthesis, pregnenolone => testosterone<br>=> dihydrotestosterone | 18.52 | 0.05 | 0.05 | 0.04 | 1 | Module |

*FDR: Fold-Enrichment; N: Gene Count; Cat: Category. All data are generated at pvalue < 0.05 and q-value < 0.05 and sorted ascendingly by p.adjust values.*

**Fig. 4:**
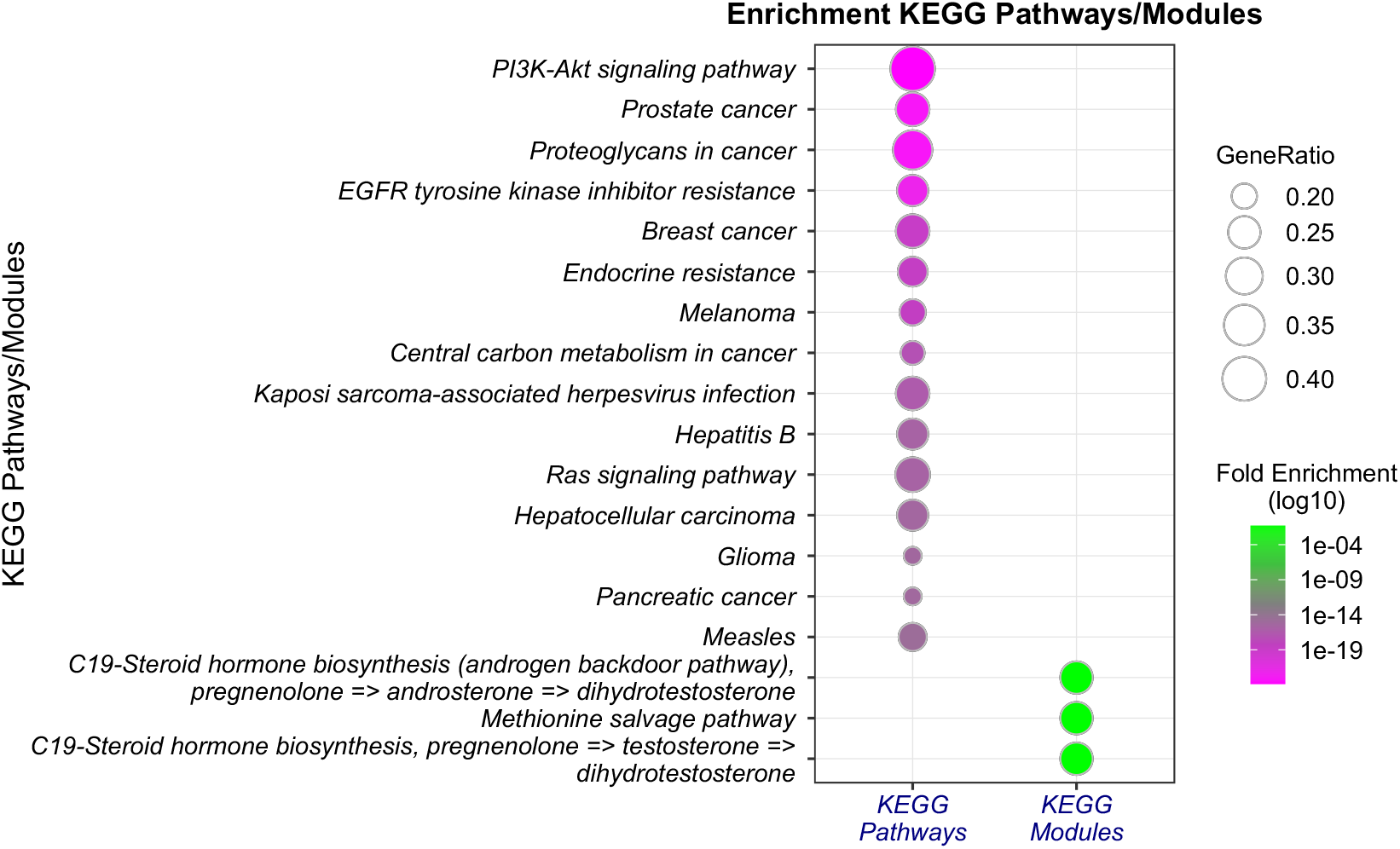
Dotplot of KEGG pathways and metabolic modules of FSK/Cancer common targets.

**Fig. 5:**
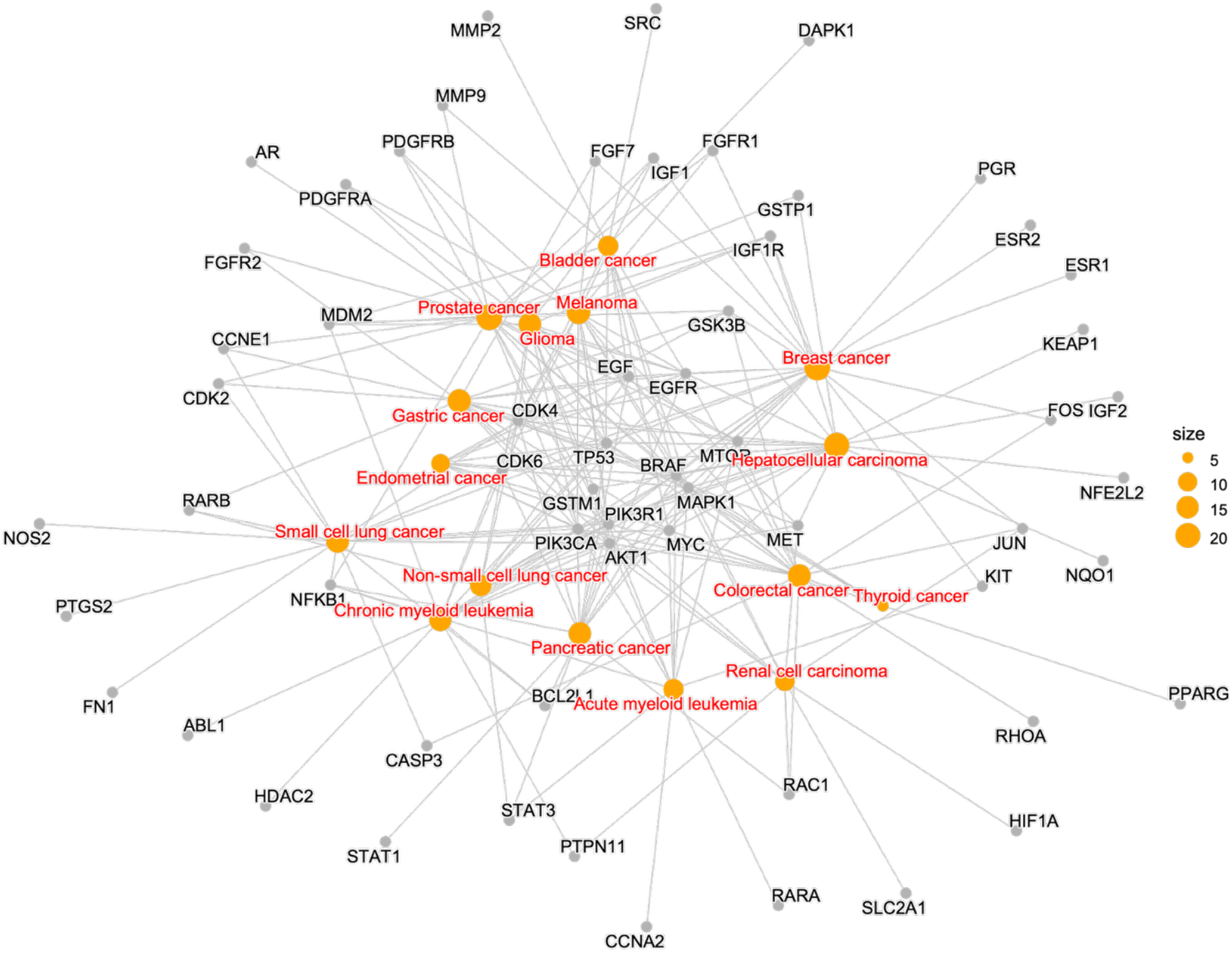
Gene-concept network graph of FSK-related cancer types and their associated genes.

**Fig. 6:**
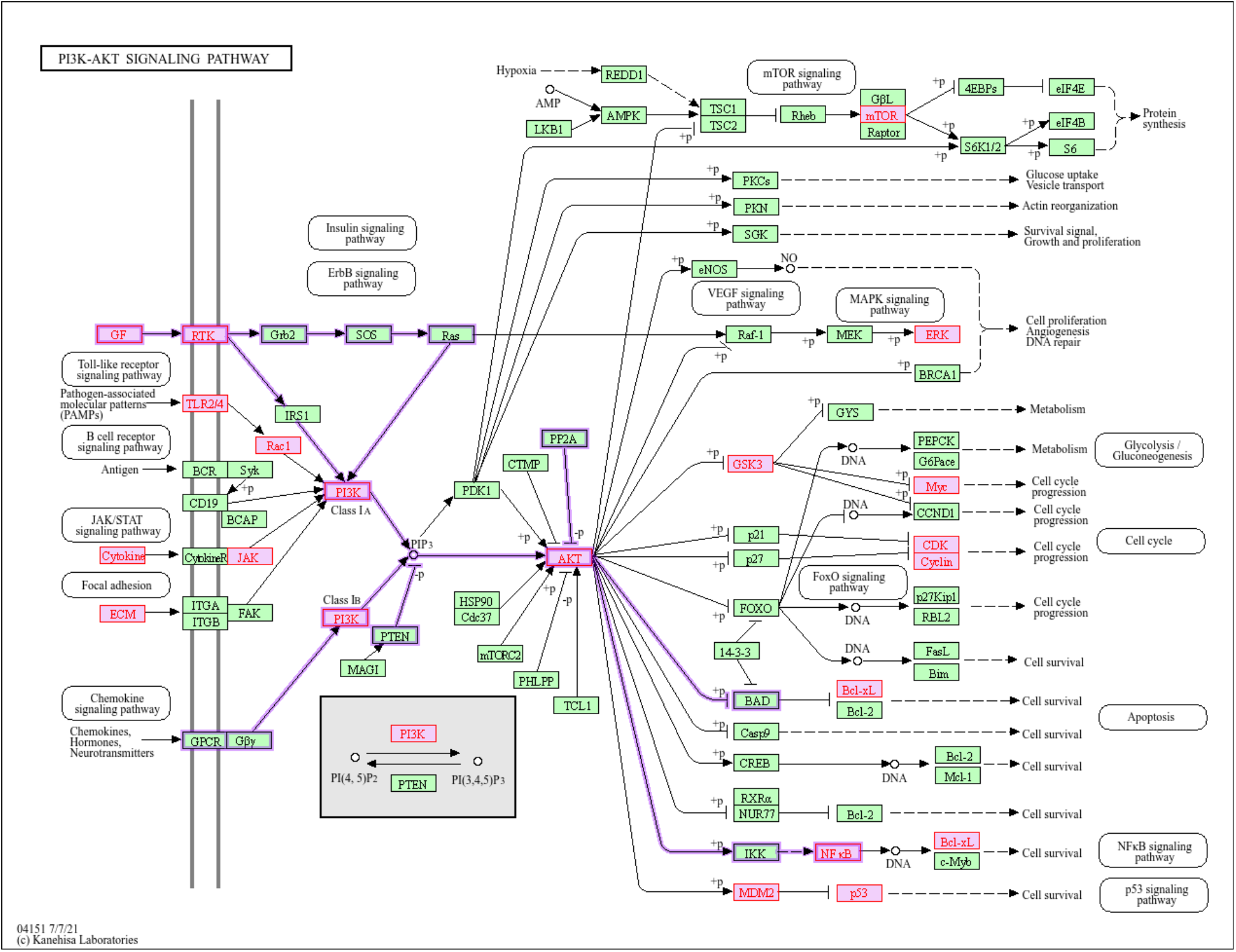
KEGG PI3K-Akt signaling pathway map. Purple highlights represent the potential core targets by which FSK would regulate cancer incidence and/or progression. Violet outlined arrows represent both PI3K (nt06530) and Akt (nt06214) oncogenic signaling pathways.

The enrichment of modules (Figure 4), including the C19-steroid hormone biosynthesis pathway (M00977 and M00976 with FDR of 22.23 and 18.52, respectively) and methionine salvage (M00034 with FDR of 18.52), aligns with the established role of PI3K/Akt in metabolic reprogramming of cancer cells (Pavlova and Thompson, 2016). Moreover, PI3K/Akt activation enhances glycolysis, lipid synthesis, and nucleotide biosynthesis, facilitating rapid tumor growth. Forskolin’s predicted impact on these modules suggests it may counteract cancer-specific metabolic adaptations.

The enrichment of FSK/Cancer targets in stress response and kinase activity GO terms (Table 1; Figure 3) suggests FSK’s dual role in mitigating oxidative stress and disrupting oncogenic signaling. Thus, its modulation of PI3K/Akt signaling, leading to apoptosis, decreased proliferation, and sensitization to chemotherapy, offers a unique oncotherapeutic for targeting cancers with high metabolic and oxidative demands.

## CONCLUSION

The present study demonstrates that forskolin’s anticancer activity is closely linked to its modulation of the PI3K/Akt signaling pathway and its associated network of oncogenic and metabolic processes. The enrichment of forskolin targets in PI3K/Akt, EGFR, and related pathways, as well as their intersection with key hub genes such as AKT1, TP53, and EGFR, highlights a multi-faceted mechanism involving inhibition of proliferation, induction of apoptosis, and disruption of cancer-specific metabolic adaptations. These findings provide a compelling molecular rationale for forskolin’s potential as a multi-targeted anticancer agent, warranting further preclinical and clinical evaluation in cancer therapy.

## REFERENCES

Bhat, S. V, Bajqwa, B.S., Dornauer, H., do Scusa, N.J., Fehlhaber, H.-W., 1977. Structures and stereochemistry of new labdane diterpiniods from coleus forskohlii briq. Tetrahedron Lett. 18, 1669–1672. 10.1016/S0040-4039(01)93245-9

Blume-Jensen, P., Hunter, T., 2001. Oncogenic kinase signalling. Nature 411, 355–365. 10.1038/35077225

Elwia, S.K., Elnoury, H.A., and Muhammad, M.H., 2018. Forskolin Effect on FOXO1 Expression and Relationship of FOXO1 Activation to Oxidative Stress: From Molecular to Therapeutic Strategy. Biomarkers J. 04, 1–7. 10.21767/2472-1646.100049

Follin-Arbelet, V., Misund, K., Hallan Naderi, E., Ugland, H., Sundan, A., Kiil Blomhoff, H., 2015. The natural compound forskolin synergizes with dexamethasone to induce cell death in myeloma cells via BIM. Sci. Rep. 5, 1–13. 10.1038/srep13001

Genovese, C., Sergio, D., Katia, M., Gianna, T., Daria, N., Salvatore, C., Franca, V., Edoardo, T., Giovanni, S., and Di Marco, R., 2018. Effects of a new combination of plant extracts plus d-mannose for the management of uncomplicated recurrent urinary tract infections. J. Chemother. 30, 107–114. 10.1080/1120009X.2017.1393587

Godard, M.P., Johnson, B.A., Richmond, S.R., 2005. Body composition and hormonal adaptations associated with forskolin consumption in overweight and obese men. Obes. Res. 13, 1335–1343. 10.1038/oby.2005.162

Hayashi, T., Hikichi, M., Yukitake, J., Wakatsuki, T., Nishio, E., Utsumi, T., Harada, N., 2018. Forskolin increases the effect of everolimus on aromatase inhibitor-resistant breast cancer cells. Oncotarget 9, 23451–23461. 10.18632/oncotarget.25217

He, L., Azizad, D., Bhat, K., Ioannidis, A., Hoffmann, C.J., Arambula, E., Eghbali, M., Bhaduri, A., Kornblum, H.I., Pajonk, F., 2025. Radiation-induced cellular plasticity primes glioblastoma for forskolin-mediated differentiation. Proc. Natl. Acad. Sci. 122, e2506044122. 10.1073/pnas.2506044122

Henderson, S., Magu, B., Rasmussen, C., Lancaster, S., Kerksick, C., Smith, P., Melton, C., Cowan, P., Greenwood, M., Earnest, C., Almada, A., Milnor, P., Magrans, T., Bowden, R., Ounpraseuth, S., Thomas, A., Kreider, R.B., 2005. Effects of Coleus Forskohlii Supplementation on Body Composition and Hematological Profiles in Mildly Overweight Women. J. Int. Soc. Sports Nutr. 2. 10.1186/1550-2783-2-2-54

Illiano, M., Conte, M., Sapio, L., Nebbioso, A., Spina, A., Altucci, L., Naviglio, S., 2018. Forskolin sensitizes human acute myeloid leukemia cells to H3K27me2/3 demethylases GSKJ4 inhibitor via Protein Kinase A. Front. Pharmacol. 9, 1–9. 10.3389/fphar.2018.00792

Li, Z., Wang, J., 2006. A forskolin derivative, FSK88, induces apoptosis in human gastric cancer BGC823 cells through caspase activation involving regulation of Bcl-2 family gene expression, dissipation of mitochondrial membrane potential and cytochrome c release. Cell Biol. Int. 30, 940–946. 10.1016/j.cellbi.2006.06.015

Liu, X., Ouyang, S., Yu, B., Liu, Y., Huang, K., Gong, J., Zheng, S., Li, Z., Li, H., Jiang, H., 2010. PharmMapper server: A web server for potential drug target identification using pharmacophore mapping approach. Nucleic Acids Res. 38, 5–7. 10.1093/nar/gkq300

Loftus, H.L., Astell, K.J., Mathai, M.L., Su, X.Q., 2015. Coleus forskohlii extract supplementation in conjunction with a hypocaloric diet reduces the risk factors of metabolic syndrome in overweight and obese subjects: A randomized controlled trial. Nutrients 7, 9508–9522. 10.3390/nu7115483

Lokesh, B., Deepa, R., Divya, K., 2018. Medicinal Coleus (Coleus forskohlii Briq): A phytochemical crop of commercial significance - Review. J. Pharmacogn. Phytochem. 7, 2856–2864.

Manning, B.D., Toker, A., 2017. AKT/PKB Signaling: Navigating the Network. Cell 169, 381–405. 10.1016/j.cell.2017.04.001

Miastkowska, M., Sikora, E., Lasoń, E., Garcia-Celma, M.J., Escribano-Ferrer, E., Solans, C., Llinas, M., 2017. Nano-emulsions as vehicles for topical delivery of forskolin. Acta Biochim. Pol. 64, 713–718. 10.18388/abp.2017_2334

Pateraki, I., Andersen-Ranberg, J., Jensen, N.B., Wubshet, S.G., Heskes, A.M., Forman, V., Hallström, B., Hamberger, Britta, Motawia, M.S., Olsen, C.E., Staerk, D., Hansen, J., Møller, B.L., Hamberger, Björn, 2017. Total biosynthesis of the cyclic AMP booster forskolin from Coleus forskohlii. Elife 6, 1–28. 10.7554/eLife.23001

Pavlova, N.N., Thompson, C.B., 2016. The Emerging Hallmarks of Cancer Metabolism. Cell Metab. 23, 27–47. 10.1016/j.cmet.2015.12.006

Pullaiah, T, 2022. Forskolin. Springer.

Pullaiah, T., 2022. Pharmacology of Coleus forskohlii and Forskolin, in: Forskolin. pp. 65–106. 10.1007/978-981-19-6521-0_5

Quinn, S.N., Graves, S.H., Dains-McGahee, C., Friedman, E.M., Hassan, H., Witkowski, P., Sabbatini, M.E., 2017. Adenylyl cyclase 3/adenylyl cyclase-associated protein 1 (CAP1) complex mediates the anti-migratory effect of forskolin in pancreatic cancer cells. Mol. Carcinog. 56, 1344–1360. 10.1002/mc.22598

Ravinder, D., Rampogu, S., Dharmapuri, G., Pasha, A., Lee, K.W., Pawar, S.C., 2022. Inhibition of DDX3 and COX-2 by forskolin and evaluation of anti-proliferative, pro-apoptotic effects on cervical cancer cells: molecular modelling and in vitro approaches. Med. Oncol. 39, 1–11. 10.1007/s12032-022-01658-3

Roshni, P.T., Rekha, P.D., 2024. Biotechnological interventions for the production of forskolin, an active compound from the medicinal plant, Coleus forskohlii. Physiol. Mol. Biol. Plants 30, 213–226. 10.1007/s12298-024-01426-9

Saleem, A.M., Dhasan, P.B., Rafiullah, M.R.M., 2006. Simple and rapid method for the isolation of forskolin from Coleus forskohlii by charcoal column chromatography. J. Chromatogr. A 1101, 313–314. 10.1016/j.chroma.2005.11.032

Salehi, B., Staniak, M., Czopek, K., Stępień, A., Dua, K., Wadhwa, R., Kumar Chellappan, D., Sytar, O., Brestic, M., Ganesh Bhat, N., Venkatesh Anil Kumar, N., del Mar Contreras, M., Sharopov, F. C., Cho, W., Sharifi-Rad, J., 2019. The Therapeutic Potential of the Labdane Diterpenoid Forskolin. Appl. Sci. 10.3390/app9194089

Sapio, L., Gallo, M., Illiano, M., Chiosi, E., Naviglio, D., Spina, A., Naviglio, S., 2017. The Natural cAMP Elevating Compound Forskolin in Cancer Therapy: Is It Time? J. Cell. Physiol. 232, 922–927. 10.1002/jcp.25650

Saxton, R.A., Sabatini, D.M., 2017. mTOR Signaling in Growth, Metabolism, and Disease. Cell 168, 960–976. 10.1016/j.cell.2017.02.004

Singh, S., Tandon, J.S., 1982. Coleonol and forskolin from Coleus forskohlii. Planta Med. 45, 62–63. 10.1055/s-2007-971249

Tripathi, C.K.M., Basu, S.K., Jain, S., Tandon, J.S., 1995. PRODUCTION OF COLEONOL (FORSKOLIN) BY ROOT uuus CELLS 0∼ PLANT COLEUS FORSKOHLII. Biotechnol. Lett. 17, 423–426.

Wang, H., Lou, C., Ma, N., 2019. Forskolin exerts anticancer roles in non-hodgkin’s lymphomas via regulating axin/ β-catenin signaling pathway. Cancer Manag. Res. 11, 1685–1696. 10.2147/CMAR.S180754

Wang, X., Pan, C., Gong, J., Liu, X., Li, H., 2016. Enhancing the Enrichment of Pharmacophore-Based Target Prediction for the Polypharmacological Profiles of Drugs. J. Chem. Inf. Model. 56, 1175–1183. 10.1021/acs.jcim.5b00690

Wang, X., Shen, Y., Wang, S., Li, S., Zhang, W., Liu, X., Lai, L., Pei, J., Li, H., 2017. PharmMapper 2017 update: A web server for potential drug target identification with a comprehensive target pharmacophore database. Nucleic Acids Res. 45, W356–W360. 10.1093/nar/gkx374

Wu, S.L., Ma, J., Qi, H.L., Zhang, Y., Zhang, X.Y., Chen, H.L., 2003. Forskolin up-regulates metastasis-related pheno-types and molecules via protein kinase B, but not PI-3K, in H7721 human hepato-carcinoma cell line. Mol. Cell. Biochem. 254, 193–202. 10.1023/A:1027392212341

